# WNK1 Enhances Migration and Invasion in Breast Cancer Models

**DOI:** 10.1101/2021.03.09.434610

**Authors:** Ankita B. Jaykumar, Jiung Jung, Pravat Parida, Tuyen T. Dang, Magdalena Grzemska, Svetlana Earnest, Chonlarat Wichaidit, Elizabeth J. Goldsmith, Gray W. Pearson, Srinivas Malladi, Melanie H. Cobb

**Author notes:** Corresponding author, 6001 Forest Park Rd., Dallas, Texas 75390-9041. Current address: Dept of Neurosurgery, Stephenson Cancer Center, University of Oklahoma Health Science Center, Oklahoma City, Oklahoma (Tuyen T. Dang).

## Abstract

Metastasis is the major cause of mortality in breast cancer patients. Many signaling pathways have been linked to cancer invasiveness, but blockade of few protein components has succeeded in reducing metastasis. Thus, identification of proteins contributing to invasion that are manipulable by small molecules may be valuable in inhibiting spread of the disease. The protein kinase WNK1 (with no lysine (K) 1) has been suggested to induce migration of cells representing a range of cancer types. Analyses of mouse models and patient data have implicated WNK1 as one of a handful of genes uniquely linked to invasive breast cancer. Here we present evidence that inhibition of WNK1 slows breast cancer metastasis. We show that depletion or inhibition of WNK1 reduces migration of several breast cancer cell lines in wound healing assays and decreases invasion in collagen matrices. Furthermore, WNK1 depletion suppresses expression of AXL, a tyrosine kinase implicated in metastasis. Finally, we demonstrate that WNK inhibition in mice attenuates tumor progression and metastatic burden. These data showing reduced migration, invasion, and metastasis upon WNK1 depletion in multiple breast cancer models suggest that WNK1 contributes to the metastatic phenotype and that WNK1 inhibition may offer a therapeutic avenue for attenuating progression of invasive breast cancers.

---

The protein kinase With No lysine (K) 1 (WNK1) is the most widely expressed of a family of four related enzymes notable for their unique catalytic lysine location, which distinguishes them from all other members of the eukaryotic protein kinase superfamily (1). Positional cloning and subsequent physiological studies demonstrated that mutations in WNK1 and WNK4 genes can cause a rare form of hypertension resulting from their increased expression (2–5). Among the best described WNK targets are the closely related protein kinases OSR1 (*OXSR1*, oxidative stress responsive 1) and SPAK (*STK39*, STE20/SPS-1-related proline-alanine-rich kinase) (6–8). These kinases form complexes with and are activated upon phosphorylation by WNKs and regulate the activities of cation chloride cotransporters that support ion homeostasis throughout the body (9–11). WNKs themselves are sensitive to changes in osmotic stress and intracellular chloride concentrations (1, 12, 13).

WNK1 is an essential gene in mouse. Knock out of the mouse WNK1 gene prevents development of a functional blood vessel architecture resulting in embryonic lethality by embryonic day 12 (14–17). Depletion of WNK1 also prevents migration and angiogenesis in assays with cultured primary human endothelial cells and prevents endothelial sprouting from aortic slices ((18), Jaykumar et al. In preparation). A similar loss of endothelial cell migration is observed upon depletion of OSR1. Although a detailed mechanistic understanding of the effects of WNK1 on endothelial migration is lacking, decreased expression of WNK1 in cultured endothelial cells caused reduced expression of a number of factors that promote angiogenesis including Slug (*SNAI2*), vascular endothelial growth factor A (VEGF-A), and matrix metalloproteinases (18). These and other findings indicate an effect of the WNK1 cascade on induction of a mesenchymal phenotype essential for endothelial wound healing (18, 19).

Aside from its important functions in vascular biology, WNK1 has been implicated in migration in multiple cancer types including glioblastoma, prostate cancer, non-small cell lung cancer, and breast cancer (20–24). In several of these studies, as well, the actions of WNK1 were associated with a shift towards a mesenchymal phenotype. We chose to focus on examining the impact of WNK1 on breast cancer migration because of an unbiased transposon insertional mutagenesis study in mice designed to identify breast cancer susceptibility genes. Using survival prediction analysis of breast cancer patient progression data, this study categorized WNK1 as one of a handful of driver genes in high-risk, invasive breast cancer (25). Here we show that in breast cancer model systems depleting or inhibiting WNK1 decreases migration and invasion in vitro. We find that WNK1 promotes invasion through a previously identified network involving Slug and the tyrosine kinase AXL (26). Finally, we show that a WNK inhibitor reduces tumor burden in a mouse xenograft model of metastatic breast cancer.

## Results

### Migration of breast cancer cells is blocked by WNK inhibition

Migration was first examined in MCF-ductal carcinoma in situ (DCIS) cells, using a wound healing assay as described (26). Following depletion of WNK1, migration of MCF-DCIS cells was reduced (**Fig 1A**). A similar extent of inhibition was also observed in cells exposed to a selective WNK inhibitor, SW133708 (**Fig 1B**), identified in a screen in the UT Southwestern High Throughput Screening Core (Supplemental Information, **Fig S1A, S1B**). Migration was decreased to nearly the same extent as that caused by cytochalasin D, an inhibitor of actin polymerization. We evaluated the requirement for WNK in migration of several breast cancer cell lines, among them 3 triple negative (TNBC) lines - BT-549, BT-20, and MDA-MB-231, and two estrogen and progesterone receptor negative, HER2 positive lines - HCC1569, and HCC1419, using the WNK inhibitor WNK463 (27). This inhibitor was shown to have high selectivity for WNKs, with remarkably little detectable inhibition of a panel of over 400 other protein kinases (28). WNK463 significantly reduced wound closure of these 5 migrating lines; it was nearly as effective as cytochalasin D for 4 of the 5 lines tested here (**Fig 1C, Fig S2**). These results indicate that interfering with WNK1, either by depletion or via two structurally distinct inhibitors, reduces migration of invasive types of breast cancer cells in two-dimensional culture.

**Figure 1.**
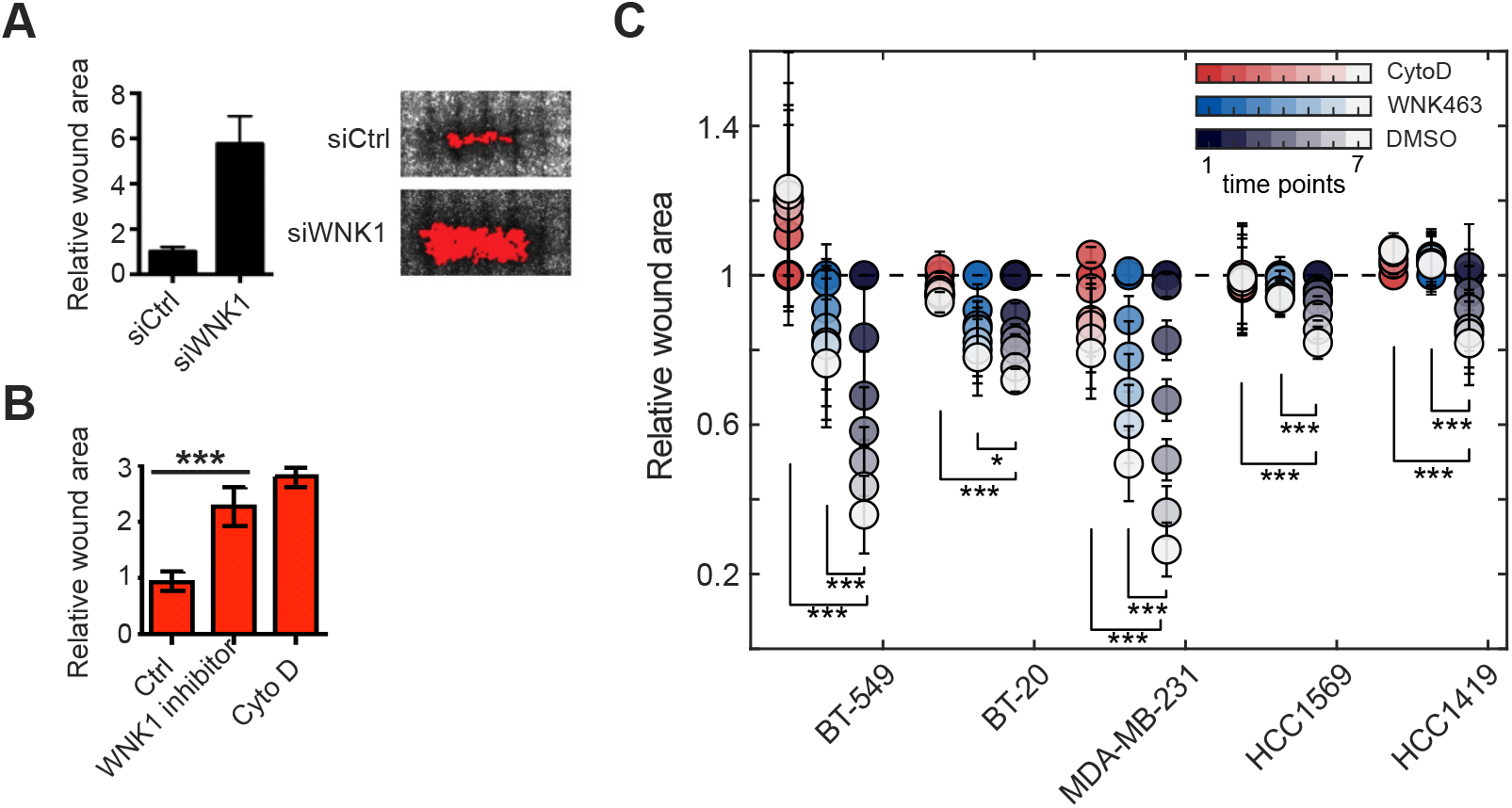
Inhibition of WNK1 decreases breast cancer cell migration. A) Wound closure of MCF-DCIS cells transfected with control or WNK1 siRNA and grown to confluence. Representative images 24 h after wounding are shown. Graph shows quantification of relative wound area (mean+/−SD, n=6 wounds). B) Wound closure of MCF-DCIS cells grown to confluence and treated with DMSO or SW133708 (10 μM) or cytochalasin D immediately after wounding. Representative images 24 h after wounding are shown. Graph shows quantitation of wound area (mean+/−SD, n=6 wounds). C) BT-549, BT-20, HCC1569 and HCC1419 cells were grown to confluence, treated with DMSO, WNK463 (1 μM) or cytochalasin D (1 μM), and imaged at time 0 and over 6 time points ending at 48 hours. Results are from n=8 and are shown as mean+/−SE. The last two time points from each treatment were used to determine statistical significance analyzed by Student’s *t*-test. * p<0.05, *** p<0.0001.

### Collagen invasion by breast cancer cells is blocked by WNK inhibition

To determine if WNK1 had an effect on the ability of cells to invade into the extracellular matrix, we examined the ability of MDA-MB-231 cells to migrate into collagen (**Fig 2**). Invasion was quantified from the number of cells migrated per field in 10 micrometer increments in 48 hours using phase contrast microscopy (29,30). Depleting WNK1 expression resulted in close to a 50% reduction in the number of invading cells per field (**Fig 2A**). We then tested the effect of the WNK inhibitor WNK463. For these experiments, 3 × 10^4^ cells were seeded on two different collagen concentrations for 24 hours prior to the addition of WNK463. Cells invading collagen were then quantified after an additional 24 hours following exposure to the inhibitor or vehicle control. With this protocol, the number of cells invading either collagen concentration was reduced by WNK463 (**Fig 2B**). At the higher collagen concentration, the number of cells invading in the presence of DMSO was smaller than at the lower collagen concentration and the reduction in invading cells due to the inhibitor was also close to 50%. At the lower collagen concentration, the inhibitor-induced decrease in invading cells was close to 20%. One factor contributing to the smaller decrease in invasion at the lower collagen concentration compared to WNK1 depletion is the 24 hours of pre-incubation on collagen prior to the addition of the inhibitor, during which time cells in both groups could invade unimpeded by the inhibitor. As was the case for migration in two dimensions, these findings suggest that WNK1 is required for invasion in a three-dimensional matrix.

**Figure 2.**
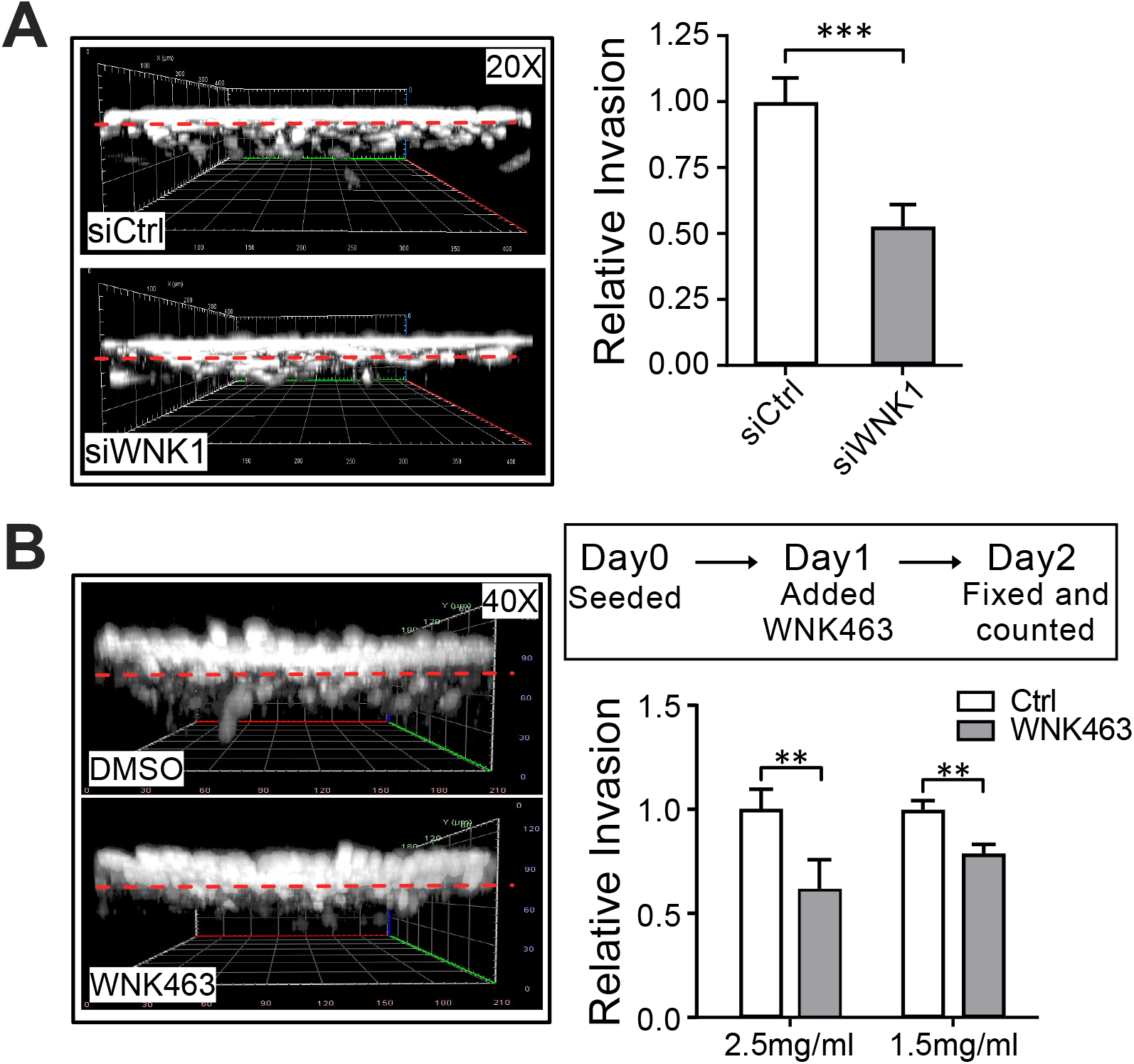
Inhibition of WNK1 decreases collagen invasion of MDA-MB-231. A. Equal numbers (1 × 10^4^) of control siRNA (siCtrl)- or WNK1 siRNA (siWNK1)-treated MDA-MB-231 cells were seeded on top of the collagen matrices (1.5 mg/ml) and allowed to invade for 48 hours at 37°C. Cells were fixed, stained and invading cells were counted from each replicate sample (n=6). The left panel shows a 3D projection reconstructed from the z-stacks of phalloidin/DAPI stained cells (grayscale images) using a confocal microscope at 20x magnification. The bar graph shows quantification of average numbers of invading cells. **B.** Twenty-four hours after seeding 3 × 10^4^ cells into 3D collagen gels (1.5 or 2.5 mg/ml), cells were treated with DMSO or WNK463 (1 μM) for 24 hours at 37°C. 3D projection image of phalloidin/DAPI stained cells (grayscale images) taken from a confocal microscope at 40x magnification. Graphs represent average numbers of invading cells (n=3). Dotted red lines indicate cell monolayer. Data are represented as mean+/−SD. ** p<0.01, *** p<0.0001, twotailed unpaired Student’s *t*-tests.

### WNK1 is in a regulatory cascade with AXL and OSR1

The Tyro3-AXL-Mer (TAM) receptor tyrosine kinase AXL was shown to form a network that promotes migration and a partial epithelial-mesenchymal transition (EMT) in MCF-DCIS cells (26,31,32). Numerous studies have suggested the importance of AXL as a target in breast and other cancers (33–36) (37–41). AXL causes feed-forward transcriptional activation of Slug (SNAI2) through autocrine transforming growth factor (TGF) β signaling as well as more directly through SMAD3 (42,43). The p53 family transcription factor p63 also regulates the expression of AXL in TGFβ-dependent EMT (44). Because we previously found interactions of WNK1 with SMAD3 function and we showed that WNK1 influences Slug expression in endothelial cells, we asked whether inhibiting WNK1 changed expression of AXL in MCF-DCIS cells (18,45). Silencing WNK1 decreased expression of AXL and Slug in this invasive MCF-10 model, as detected by immunoblotting, similar to the known AXL regulator p63 (**Fig 3A**). A significant reduction of AXL expression upon WNK1 depletion was observed in MDA-MB-231 cells (**Fig 3B, E, F**) and a smaller but clear reduction was noted in the triple negative breast cancer line SUM159 (**Fig 3C**). In contrast, WNK1 depletion did not impair expression of its substrate kinase OSR1, although as much as half of WNK1 and OSR1 are co-localized in a number of cell types (46) (**Fig 3F**). Similar to WNK1 siRNA, the WNK inhibitor WNK463 decreased expression of AXL in MDA-MB-231 cells (**Fig 3D**). Confirmation of the effect of WNK1 depletion on AXL expression was further demonstrated by immunofluorescence in MDA-MB-231 cells (**Fig 3G, H**).

**Figure 3.**
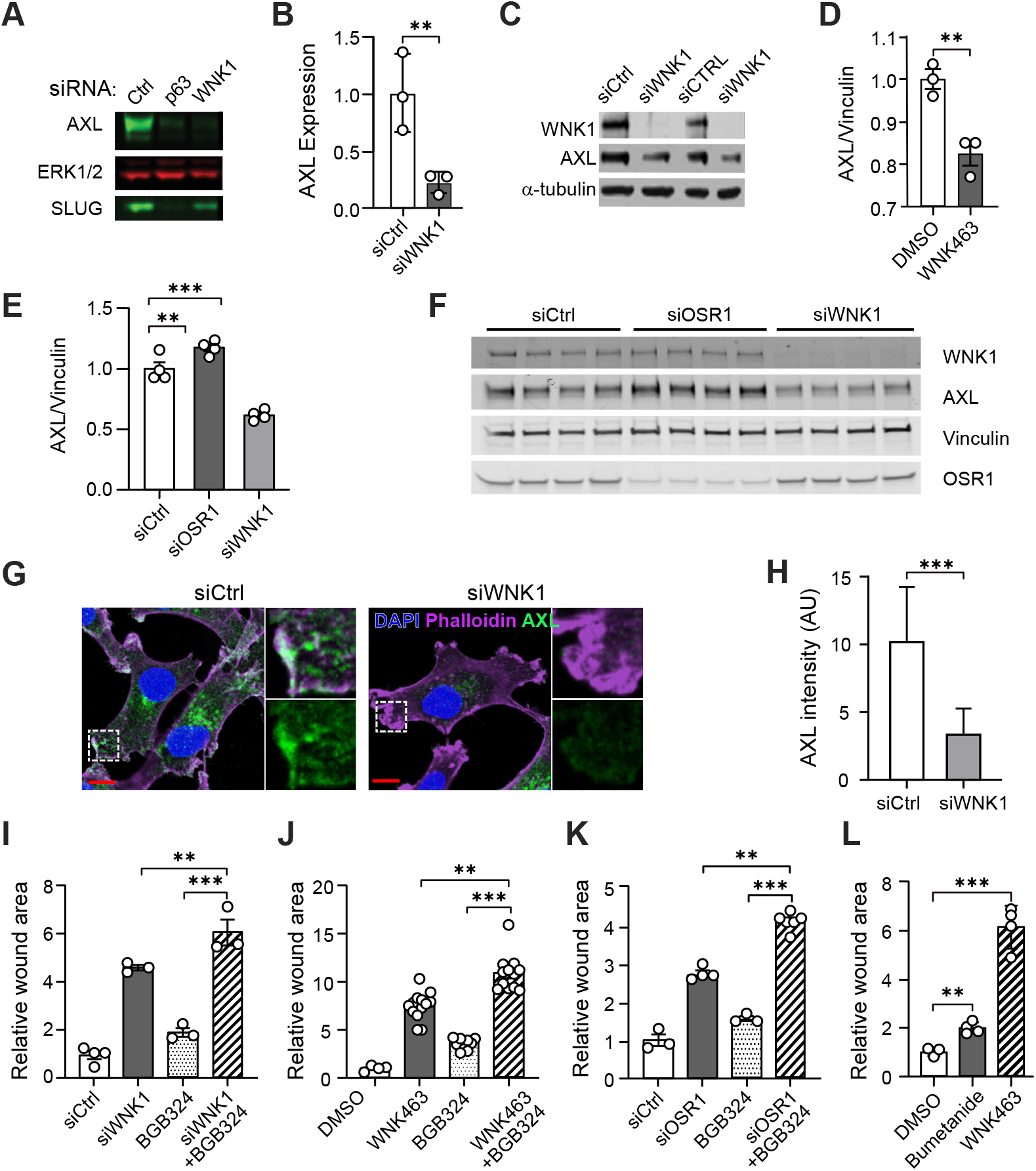
Inhibition of WNK1 reduces Axl expression. **A**) MCF-DCIS cells (70% confluent) were treated with either control, p63 or WNK1 siRNA for 72 h. Cells were lysed and immunoblotted for AXL, and Slug, ERK as the loading control. **B**) Quantification of normalized AXL expression from A; n=3. **C**) SUM159 cells treated with siCtrl or siWNK1 were harvested 72 h after transfection and lysates were analyzed by immunoblotting with the indicated antibodies. α-tubulin is the loading control. **D**) Quantification of Axl protein expression from MDA-MB-231 cells treated with WNK463 (1 μM) overnight (n=3). **E**) Quantification of AXL protein expression from MDA-MB-231 cells treated with control, OSR1 or WNK1 siRNA overnight (n=4). **F**) Immunoblots from E. **G**) MDA-MB-231 cells were treated with WNK1 or control siRNAs for 72 hr. Cells were stained for AXL (green), F-actin (phalloidin, purple), and nuclei (DAPI, blue). Representative images of AXL immunofluorescence staining. Areas marked with white squares are amplified and shown in insets. Scale bars 10μm. **H**) Quantification of mean AXL immunofluorescence intensity as shown in G. An unpaired t-test was performed. *** p < 0.0001, N = 60. AU, arbitrary units. Panels **I, J, K, L**: Migration of MDA-MB-231 cells treated as indicated with wound area normalized to that at time 0. **I**) Control siRNA (n=4), WNK1 siRNA (n=3), BGB324 (2 μM) (n=3), and WNK1 siRNA+BGB324 (n=3) for 24 hours. **J**) DMSO (n=4), WNK463 (1 μM) (n=13), BGB324 (2 μM) (n=10) and WNK463+BGB324 (n=12) for 36 hours. **K**) Control siRNA (n=3), OSR1 siRNA (n=4), BGB324 (2 μM) (n=3), and OSR1 siRNA+BGB324 (n=6) for 24 hours. **L**) DMSO, bumetanide (20 μM) and WNK463 (1 μM) overnight (n=4 for each). Data in panels **I, J, K**, and **L** are mean+/−SE; analyzed by 1-way-ANOVA. **p<0.01, ***p<0.0001.

Disruption of the OSR1 gene in endothelial cells in mouse recapitulated the WNK1 gene disruption phenotype (16). As a result, we asked if OSR1 depletion also reduced AXL expression in breast cancer cells. In contrast to WNK1 siRNA, OSR1 siRNA did not decrease AXL protein (**Fig 3E,F**); in fact, a small increase in AXL was detected in OSR1-depleted cells. We then tested effects of the AXL inhibitor BGB324 on migration to determine the relative ability of each component to regulate migration. The AXL inhibitor BGB324 alone reduced migration of MDA-MB-231 cells, as previously reported (47), but its effects were less pronounced than depletion of WNK1 (**Fig 3I**), the inhibitor WNK463 (**Fig 3J**), or depletion of OSR1 (**Fig 3K**). Nevertheless, modest but significant effects of BGB324 could be detected even in cells in which WNK1 or OSR1 had been depleted as well as in WNK463-treated cells, suggesting that their effects may be additive. From these findings, it appears that the control of AXL expression by WNK1 occurs through a mechanism independent of its effector kinase OSR1. The results also suggest that control of AXL expression alone is not sufficient to account for the actions of WNK1 on migration.

OSR1 knockdown in cell culture prevented endothelial cell migration (18). As noted above, depletion of OSR1 also attenuated migration of MDA-MB-231 cells (**Fig 3K**), consistent with its actions in endothelial cells. Among best validated targets of OSR1 are the sodium, potassium, two chloride cotransporters, NKCC1 and NKCC2, that regulate ion balance and cell volume (9-11). NKCC1 is broadly expressed in tissues and has been implicated in the invasiveness of certain cancers such as glioblastoma (20,48). NKCCs are inhibited by furosamide-related diuretics such as bumetanide. We compared effects of bumetanide to WNK463 and observed that bumetanide inhibited migration of MDA-MB-231 cells, but not nearly as effectively as WNK463 (**Fig 3L**). These findings suggest that OSR1 is a significant mediator of the migratory ability of WNK1; however, NKCC1 is only one of multiple inputs that contribute to WNK1-dependent migration downstream of OSR1.

### WNK1 activity is elevated in cells isolated from breast cancer metastases

A study by Kang et al (49) examined features of breast cancer cells that colonized bone by injecting MDA-MB-231 cells into mice, and then re-isolating and analyzing properties of the cells with a propensity to metastasize to this site. We used MDA-MB-231 parental cells and bone metastatic derivatives (MDA-BOM) isolated as previously described to evaluate any differences in WNK activity in these cells (49). Because OSR1 is phosphorylated and activated by WNK, OSR1 phosphorylation, detected with phospho-antibodies to activating phosphorylation sites, is a useful readout of WNK activity. To estimate relative WNK activity in the parental cells and cells derived from bone metastases, lysates of these cells were immunoblotted for phospho-OSR1. The amount of phospho-OSR1 relative to total OSR1 in MDA-MB-231 BOM cells was nearly twice that in the parental MDA-MB-231 cells (**Fig 4A,B**).

**Figure 4.**
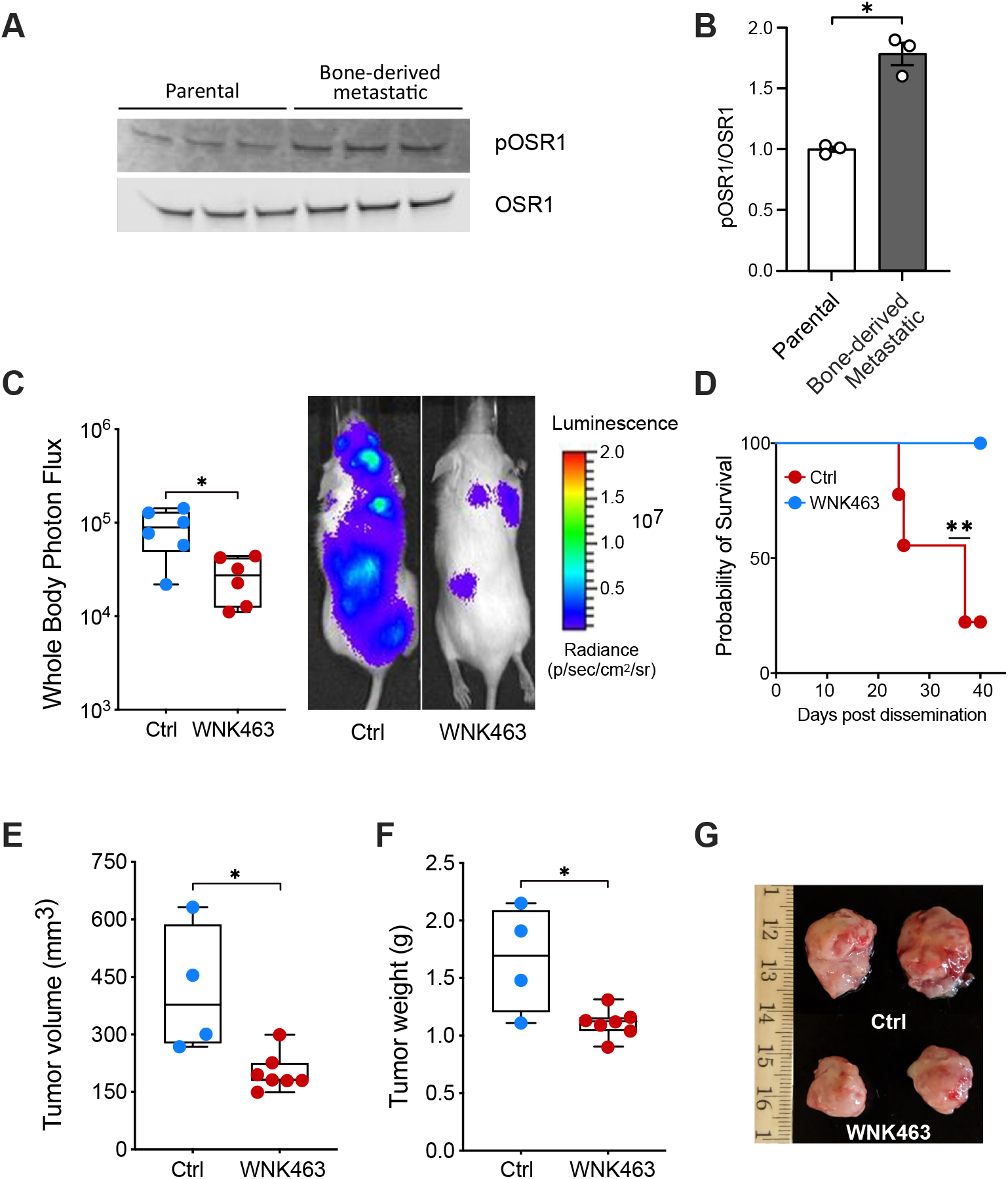
The WNK inhibitor WNK463 decreases tumor burden in mice. **A.** Lysates from MDA-MB-231 parental cells and bone metastatic cells were immunoblotted with OSR1 and pOSR1 (pT325) antibodies. **B.** Quantitation of blots in A; *p<0.05. **C.** 10^5^ MDA-MB-231 cells transduced with GFP-luciferase (MDA231-BoM) were injected intracardially into 5-6 week-old NSG (*NOD.Cg-Prkdcscid Il2rgtm1Wjl/SzJ*) mice (Jackson Labs). WNK463 or vehicle was administered by oral gavage every other day, three-days post injection, at 1 mg/kg for week 1, and at 1.5 mg/kg for weeks 2-4. Mice were injected with D-luciferin (150 mg/kg) retro-orbitally and imaged non-invasively using an IVIS spectrum instrument (PerkinElmer) weekly. Wholebody photon flux is plotted on the left, *p<0.05. Representative mice receiving vehicle or WNK463 are shown on the right. **D.** Survival is plotted; **p=0.01. **E.** 10^5^ MDA-MB-231 bone metastatic cells transduced with GFP-luciferase as in C. above were mixed with matrigel and injected orthotopically in 5-6 week old NSG mice. Two-weeks post injection, mice with palpable tumors were treated with either control or WNK463 by oral gavage every other day. During week 1 the dose was 1 mg/kg, and increased from weeks 2-4 to 1.5 mg/kg. Tumor volume for control and WNK463-treated mice was estimated from caliper measurements; *p=0.05. Student’s *t* test (unpaired). **F.** Weights of tumors removed from control and WNK463-treated mice; *p=0.05. **G.** Representative tumors for each treatment are shown.

### Tumor volume is reduced in mice injected with bone metastatic cells and treated with WNK463

Based on the evidence above that WNK1 inhibition reduces migration and invasion of breast cancer cells, we determined the effectiveness of WNK463 in blocking tumor progression and metastasis in mice using MDA-MB-231 xenograft models. Because increased phospho-OSR1 suggested higher WNK activity in MDA-MB-231 BOM cells, we injected these cells intracardially into 5-6 week-old immune compromised (NSG) mice to examine effects of WNK inhibition on metastasis. WNK463 was shown to reduce hypertension in mouse models, but was associated with toxicity at the most effective concentrations reported (27). Because of these concerns for in vivo use, we chose relatively low doses (1-1.5 mg/kg) of drug rather than higher doses tested for hypertension (up to 10 mg/kg (27)). At these lower doses, WNK463 substantially decreased overall metastatic burden, as assessed by bioluminescence (**Fig 4C**). In addition, treatment with WNK463 was associated with improved survival compared to the vehicle-treated animals (**Fig 4D**). To test the potential efficacy of WNK463 on the growth of orthotopic tumors, MDA-MB-231 cells were injected into mammary fat pads. WNK463 or DMSO was administered for 4 weeks after tumors were palpable (**Fig 4E-G**). Using this protocol, tumors in the inhibitor-treated animals were smaller by volume and weight than those in the control animals. These results provide evidence that WNK1 inhibition attenuates tumor progression and metastasis. These findings indicate that WNK1 may be a worthwhile target to be considered for therapy of invasive breast cancer.

## Discussion

Invasive breast cancer is the second leading causing of mortality in women (50). Poor survival is associated with metastasis to distant sites including bone, brain, liver, and lung (51). The majority of therapeutic approaches for breast and other cancers have focused on cell proliferation, while success in interfering with the migratory and invasive events that reduce survival of breast cancer patients has been more limited (32). This problem motivated our study to evaluate WNK1 as an initiator of invasion.

WNK1 was recognized early as a regulator of blood pressure (2). Studies in mice revealed that, in addition to well-documented actions on ion transport, WNK1 is required for the elaboration of the circulatory system through its essential function in endothelial angiogenesis (14–17). Paralleling these findings in endothelial cells, WNK1 has been implicated in migration with EMT features through knockdown studies in other cancer types, as well as a contributor to stem-like properties in metastatic breast cancers (22–24). Mechanistic information has suggested links to expression of EMT factors Slug and Snail, microRNA networks, changes in expression of cell surface proteins, altered vesicle trafficking, and effects on actin polymerization (22,23,52). Genetic analysis integrating data from insertional mutagenesis and human patients has further supported the idea that WNK1 may be a relevant sensitivity in invasive breast cancer (25). In testing this concept, we find that interfering with WNK1 reduced the migratory behavior of breast cancer cells in two- and three-dimensions in vitro, reduced the dispersion of metastatic cells in mice, and reduced orthostatic tumor volume. In addition to elevated phospho-OSR1 in bone metastatic cells, we also found that lung-derived metastatic MDA-MB-231 had higher phospho-OSR1 (data not shown), suggesting that increased WNK activity may be a feature of breast cancers that can metastasize to multiple sites.

Previous findings on WNK1 in endothelial migration indicated a large number of factors that may contribute to WNK1-dependent migration, some or all of which may be relevant in breast cancers (18). Top among these is OSR1, which in contrast to many other WNK1 partners, is a direct WNK1 substrate. Disruption of the OSR1 gene in endothelial cells in mouse recapitulated the impaired angiogenesis caused by WNK1 gene disruption (16). Here we show that depletion of OSR1 also reduced migration of breast cancer cells. Inhibition of the OSR1 substrate NKCC has been shown to reduce migration in glioma (20,48); thus, we examined its contribution to breast cancer migration and found that its inhibition was not as effective as OSR1 depletion in slowing migration of breast cells. As a result, we expect that other events downstream of OSR1, in addition to this cotransporter, also contribute to the breast cancer migratory phenotype.

Surprisingly, we found that inhibiting WNK1 decreased expression of the tyrosine kinase AXL. Preliminary results suggest that this is not due to a change in AXL mRNA. This effect of WNK1 was independent of OSR1. AXL expression is associated with metastasis and poor prognosis in a variety of tumor types including breast cancer (40,47,53,54). BGB324 is a first-in-class AXL inhibitor, currently in phase II clinical trials, exhibiting promising therapeutic characteristics. It displays AXL-selective anti-tumor and anti-metastatic activity in murine models of breast cancer (47). The connection revealed here between AXL and WNK1 raises the possibility that WNK1 may be a therapeutic option in other AXL-dependent tumor types as well. AXL inhibition in tumor cells decreases the secretion of pro-angiogenic factors such as endothelin and VEGF-A and impairs functional properties of endothelial cells in vivo, suggesting its important role in the initiation of tumor angiogenesis (54). Interestingly, we previously reported that mRNAs encoding these factors were also decreased upon knockdown of WNK1 in endothelial cells (18). Perhaps WNK1 knockdown diminishes these mRNAs in part via suppression of AXL (55). We also found that inhibition of WNK1 had a more pronounced effect than the AXL inhibitor to attenuate migration in MDA-MB-231 cells. Yet, the combination of AXL and WNK inhibitors was more effective at reducing migration than inhibition of either alone. This observation warrants future studies examining the combination of AXL and WNK inhibitors on tumor progression and metastasis in animal models, which could potentially inform future clinical trials.

WNK463 is the first reported highly selective WNK inhibitor. It displays 100% oral bioavailability in C57BL/6 mice. At a dose of 10 mg/kg, WNK463 decreased blood pressure and produced multiple other clinical signs, consistent with the broad physiological actions of WNKs (27). Given the high WNK selectivity of the molecule, it remains to be determined how many of the observed clinical changes were the result of on-target actions of the inhibitor or instead may have been due to off-target effects on molecules outside the kinase family. However, in this study, not only did the inhibitor decrease tumor size and metastatic burden at the lower doses we used (1-1.5 mg/kg), no obvious untoward effects of the inhibitor were observed. No drug-treated animals died over the 4-week time course and body weights of the treated mice were not different from the control mice. These results offer promise that there will be a therapeutic window for WNK inhibitors to treat metastatic breast cancer.

## Methods

### Reagents and consumables

Reagents were obtained from the following sources: WNK463 and BGB324 (Selleck Chemicals, S8358 and S2841), anti-Axl antibody C89E7 (Cell Signaling, 8661S), anti-Vinculin antibody (Sigma Aldrich, B9131), anti-pOSR1 antibody (EMD Millipore, 07-2273), anti-OSR1 (Cell Signaling, 3729S), anti-WNK1 antibody (Cell Signaling, 4979S), anti-p63 (Cell Signaling, 4892), Optimem (Invitrogen, 51985-034), 2-hydroxypropyl-beta-cyclodextrin (Sigma Aldrich, H107-5G), Pluronic F68 (BASF corporation), cytochalasin D (Millipore Sigma, C2618), bumetanide (Sigma Aldrich, B3023), 96-well plates (Corning, 3904 or Greiner, 655090); Alexa Fluor 647 and DAPI Fluoromount-G (00-4959-52) from Thermo Fisher Scientific; phalloidin (Invitrogen, A22283 or Life Technologies, A22287); Hoechst (Invitrogen, H1399); α-tubulin, Developmental Studies Hybridoma Bank 12G10.

### Small-interfering RNAs (siRNAs)

Oligonucleotides encoding siRNA for human WNK1 (siWNK1: 5’ CAGACAGUGCAGUAUUCACTT 3’) and control siRNA (#4427037) as in (56) and OSR1 siRNA (s19303 Silencer^®^ Select) were purchased from Thermo Fisher Scientific. Transfections of siRNA were performed using Lipofectamine RNAiMAX (ThermoFisher Scientific, #13778150) according to the manufacturer’s protocol. All siRNAs were used at a final concentration of 20 nM, except in MCF-DCIS as noted.

### Cell culture and treatment

MCF-DCIS cells were as described (26). MDA-MB-231 cells (from ATCC or the Whitehurst lab, UT Southwestern) were maintained in Dulbecco’s modified Eagle’s medium (DMEM, Sigma-Aldrich, D5796) with 10% fetal bovine serum (FBS, Sigma-Aldrich, F0926) and 1% penicillin/streptomycin (Thermo Fisher Scientific, SV30010). SUM159 (Asterand) cells were cultured in Ham’s F-12 medium (Sigma-Aldrich, N6658) with 5% FBS, 5 μg/ml insulin (Sigma-Aldrich, I0516), and 1 μg/ml hydrocortisone (Sigma-Aldrich, H0888). BT-549, BT-20, HCC1569, and HCC1419 lines were obtained from the Whitehurst laboratory, UTSW, and grown in RPMI-1640 (Sigma-Aldrich, R8758) with 5% FBS and 1% penicillin-streptomycin. Bone-derived metastatic (BOM) MDA-MB-231 cells were obtained from parental MDA-MB-231 as described (49). Cell lines were either fingerprinted (PowerPlex 1.2 Kit, Promega) or independently authenticated annually (ATCC, IDEXX Bioresearch, ME) and mycoplasma-free (e-Myco Kit, Boca Scientific or Universal Mycoplasma Detection Kit 30-1012K, ATCC). All cells were maintained at 37°C and 5% CO_2_. For immunoblotting, cells were grown to 60% confluency and treated with the indicated siRNA (OSR1 or WNK1) overnight and lysed before cells attained 75% confluency for immunoblotting. Cells were treated with inhibitors WNK463 or BGB324 overnight prior to lysing and immunoblotting.

### Wound healing assays

Depletion and drug treatment experiments in MCF-DCIS cells were performed as described in (26). Briefly for depletion, 10,000 cells per well were reverse transfected with 50 nM siRNA in 96-well plates and grown to confluence for 72 h prior to wounding using a 96-pin wounding tool (V–P Scientific, FP6-WP). For inhibitor experiments, cells were grown and wounded as above. In all assays MCF-DCIS cells were fixed and stained with phalloidin and Hoechst. Montage images (5 × 4) of cells were acquired on a BD Pathway 855 microscope using a 10x objective (Olympus, UplanSApo 10x/0.40). The empty space in the well was defined with Pipeline Pilot software using a custom analysis protocol that determines empty space via a threshold of pixel intensity. Wound closure rate is inversely proportional to empty space. For depletion experiments in MDA-MB-231 cells, 80% confluent cells were treated with siRNA overnight. Wounds were made after 24-48 hours, once the cells attained 100% confluence. The cells were imaged with a Zeiss Axio Zoom V16 microscope at time 0 and 24-36 hours after treatments until the wound area in the DMSO condition was >90% closed. For inhibitor assays, the medium containing inhibitors or vehicle were replenished every 24 hours. For high-throughput wound healing assays (Figure 1), cell lines were seeded on 96-well plates. The following day, wounds were made using the pin tool above. Cell debris was removed with an EL406 plate washer (BioTek, Winooski, VT) and fresh medium was added. Compounds were previously dissolved in anhydrous DMSO. After wound creation, an Echo555 (Labcyte, San Jose, CA) acoustic liquid dispenser was used to add compound to the wounds at a final DMSO concentration of 0.1%. All plates included replicate wells with 0.1% DMSO as a neutral control. Plates were returned to a Cytomat 2C automated incubator (ThermoFisher) immediately after addition. Plates were imaged using an IN Cell Analyzer 6000 automated microscope (GE Healthcare) with a heated stage using a 10X objective and transmitted light. 12 overlapping fields centered around the monolayer wound were captured for each well. The 12 fields were then stitched together into a single image post-acquisition using the GE Developer Toolbox (v1.9.3; GE Healthcare). All plates were imaged at 8-hour intervals for a total of 7 time points, with the first time point acquired immediately after compound addition. The wound area was measured using ImageJ software (57). The remaining open area was then normalized to that in the DMSO control at 0 time. The lower the ratio (less than 1) means the wound area was closing. Cytochalasin D was used as a positive control.

### Three-dimensional (3D) collagen invasion assays

Invasion assays were adapted from Jung et al. (29). Briefly, collagen type I (Corning, #354249) was diluted to 1.5 or 2.5 mg/ml in DMEM medium. Thirty μl of the unsolidified collagen mixture was placed on the bottom of 96-well plates and solidified for 45 min at 37°C. siCtrl- or siWNK1-treated MDA-MB-231 cells were seeded at a density of 1 - 3 × 10^4^ cells per well in 0.1 ml of medium, and the 3D collagen gel culture was incubated at 37 °C. After 48 hours, cells were fixed in 3% glutaraldehyde and stained with 0.1% toluidine blue. For inhibitor studies, cells were treated with 1 μM WNK463 or DMSO as the vehicle control at 24 hours after seeding. The following day, cells were fixed and stained as described above. Invasion density was quantified by counting cells below the plane of the monolayer in five random microscopic fields per well under a light microscope (SMZ645, Nikon, Japan) at 20x magnification. Representative 3D images were obtained using a Zeiss LSM880 inverted confocal microscope (Carl Zeiss, Oberkochen, Germany).

### Immunofluorescence staining

MDA-MB-231 cells were transfected with control siRNA or WNK1 siRNA. After 36 hours, cells were seeded in 8-well chamber slides (Ibidi, #80827, Munich, Germany) at densities of 2 × 10^4^ cells/well and incubated for an additional 36 hours. Cells were fixed with 4 % paraformaldehyde and permeabilized for 20 minutes with 0.3 % Triton X-100 in PBS and blocked for 30 min at room temperature with blocking solution (5 % normal goat serum / 0.1% Triton X-100 / 1X PBS). Cells were incubated with the primary AXL antibody diluted in 1X PBS with 5 % BSA overnight at 4 °C. Subsequently, cells were incubated with an Alexa Fluor^®^ 488 conjugated goat-anti-rabbit secondary antibody (Invitrogen, A-11008) and Alexa Fluor^®^ 647 Phalloidin for 1 h at room temperature, and the slides were mounted with DAPI Fluoromount-G. Immunofluorescence images were acquired using a Zeiss LSM880 inverted confocal microscope and the fluorescence intensity of each cell was analyzed using ImageJ (NIH).

### Mouse studies

All animal studies were performed according to UTSW Institutional Animal Care and Use Committee guidelines (Animal protocol number #2017-102099). 4-5 week-old NSG mice were purchased from the UTSW animal core. For orthotropic implantation assays, 2.5 × 10^6^ MDA-MB-231 BOM cells were resuspended in 1 ml PBS and Matrigel (1:1). Mice were anesthetized by controlled isoflurane administration through a nose cone in a sterile hood. An incision was made between the fourth and fifth nipple of the mouse to expose the mammary fat pad, and 0.1 ml cell suspension was injected using a 28 gauge insulin syringe. Mice were weighed (~25g) and orally gavaged daily with 0.25 ml containing 25 μg of WNK463 at 1 or 1.5 mg/kg, as indicated, formulated as a suspension in 0.5:0.5:99 (w:w:w) 2-hydroxypropyl β-cyclodextrin:Pluronic F68:purified water. Tumor progression was monitored by measuring the volume of the tumor with Vernier calipers every week. Tumor volume was calculated by using the formula *V* = 0.5 × *a* × *b^2^*, where “a” and “b” indicate major and minor diameter. At the end point, surgically resected tumors were weighed.

For metastatic colonization assays, 1.0 × 10^5^ MDA-MB-231 BOM cells resuspended in 0.1 ml PBS were intracardially injected into the right ventricle of mice with a 26 gauge tuberculin syringe. Animals were treated daily with WNK463 as above. Tumor progression and metastatic incidence was tracked weekly by bioluminescent imaging (BLI) using an IVIS (In Vivo Imaging System) Spectrum (PerkinElmer).

Inclusion and exclusion criteria: All animals were included in the data analysis; outliers were not excluded. Attrition: Animal attrition over time due to metastasis is documented in Fig. 4D. No attrition was observed in mice bearing orthotopic tumors. Sex as a biological variable: Because this is a breast cancer study, we used only female mice for experimental analysis. Randomization: For inhibitor treatment, mice were randomized post-injection based on bioluminescent (BLI) signal. Blinding: not applicable. Power analysis: Based on previous experience with these models, animal cohorts of 5-10 mice per experimental condition are sufficient to detect differences between groups with 90% power and a 5% type I error rate. Replication: Representative figure showing one of two animal experiments.

### Statistics

Results are expressed as mean ± SEM. For comparison between only two groups two-tailed student’s t test was used. Single intergroup comparisons between 2 groups were performed with a 2-tailed one-way ANOVA. Two-way ANOVA was used to determine differences between means in two treatments and groups. p < 0.05 was considered statistically significant.

## Acknowledgments

The authors thank Mike Kalwat (Indiana Biomedical Research Institute) and members of contributing labs for valuable suggestions, Matt Esparza for assistance with MCF-DCIS cells, and Dionne Ware for administrative assistance. These studies were supported by NIH R01 HL147661 and Mary Kay Foundation grant 18-18 to MHC, Cancer Prevention and Research Institute of Texas (CPRIT) grant RP190421 and NIH R01 DK110538 to EJG, NCI R01 CA155241 and R01 CA218670 to GWP, CPRIT RP170003 to SM, American Heart Association postdoctoral fellowship 18POST34030438 to ABJ, CPRIT training grant RP160157 for support of PP, MG and early support of ABJ, Welch Foundation grants I11243 and I1128 to MHC and EJG, respectively, and the Simmons Comprehensive Cancer Center NCI grant (P30 CA142543) for assistance from the UTSW High Throughput Screening (HTS), Small Animal Imaging, and Microscopy Cores. High throughput wound healing assays (Fig. 1) were performed in collaboration with Hanspeter Niederstrasser, Ph.D., in the High Throughput Screening Core. This work was also in part supported by an NIH-sponsored S10 grant (1S10OD018005-01 to Bruce A. Posner) for the IN Cell Analyzer 6000.

## Notes

The authors declare no conflicts of interest.

### Competing Interest Statement

The authors have declared no competing interest.

