## Supplementary Figure 1 and 2 for "WNK1 Enhances Migration and Invasion in Breast Cancer Models"

### Figure S1

**A**

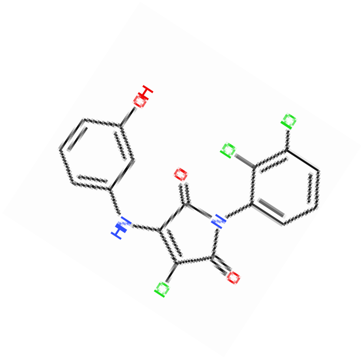


**B**

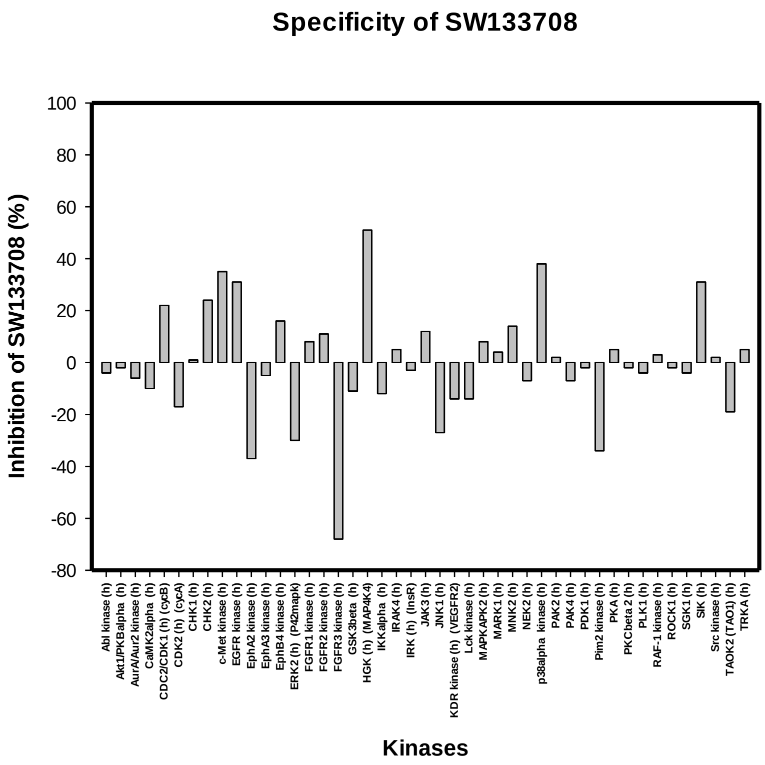


**Supplementary Figure 1. Identification of a WNK1 inhibitor from the High Throughput Screening Core. A.** A 3500 compound library, chosen by Bruce Posner, Director of the HTS core, to be representative of the UT Southwestern 230,000 compound library chemical space was screened for inhibitors of active WNK1 194-483 prepared as described (X Min et al, Structure 12, 1303, 2004) using myelin basic protein as substrate with protocols generally as described in (Piala et al, Bioorg Med Chem Lett 26, 3923, 2016). Compound SW133708 (Chembridge compound #6505063) was identified as the top hit from the screen with an $IC_{50}$ of 5 µM. **B.** A 45 human kinase panel was evaluated for inhibition by 10 µM SW133708 by the Eurofins commercial screening service (Celle-L'Evescault, France) which used HTRF (Homogeneous Time Resolved Fluorescence) to determine relative inhibition. Inhibition higher than 50% was considered significant. In addition to inhibiting WNK1, SW133708 shows inhibitory activity only toward fibroblast growth factor receptor-3 (FGFR3) of the kinases in the specificity screen.

**Figure S2**

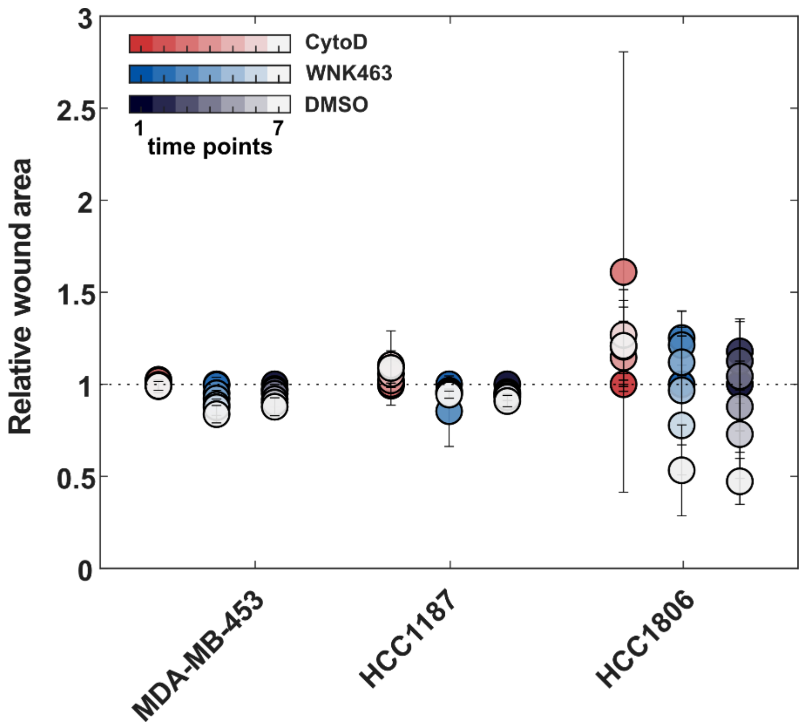


**Supplementary Figure 2. Migration assays with additional breast cell lines.** Additional cell lines were tested using the same protocol as in main Figure 1C. These lines showed little or no migration over the 48-hour time course**.**
